# COVID-19 and Active Tuberculosis Exhibit Distinct Maturation-Associated Neutrophil Transcriptional Programs

**DOI:** 10.64898/2026.11.11.765861

**Authors:** Samuel G. Harrington, Apoorv Shanker, Jessica Z. Kubicek-Sutherland

**Affiliations:** Physical Chemistry and Applied Spectroscopy, Chemistry Division, Los Alamos National Laboratory, Los Alamos, NM, United States

**Author notes:** Corresponding author (JKS).

**Keywords:** Neutrophils, RNA-Seq Analysis, COVID-19, Tuberculosis, Neutrophil Maturation, Degranulation, Chemotaxis, NETosis

## Abstract

Neutrophils can promote pathogen control or drive tissue-damaging inflammation, but the mechanisms that determine these divergent outcomes remain poorly defined. COVID-19 and active tuberculosis (TB) offer a purposeful comparison since both are pulmonary infections with substantial neutrophil involvement, yet differ in pathogen class, inflammatory duration, and the balance between neutrophil-associated host defense and pathology. We therefore tested whether their divergent neutrophil programs align with distinct maturation-associated transcriptional states. We integrated six publicly available whole-blood bulk RNA-sequencing datasets, comprising three COVID-19 and three active TB cohorts. A curated panel of 770 genes covered activation, degranulation, neutrophil extracellular trap (NET) formation, chemotaxis, cell surface receptors, and maturation. Study-associated variation was reduced using ComBat-Seq, and differential expression relative to disease-appropriate controls was assessed with DESeq2. Most genes changed in the same direction in both diseases, but 348 (45.2%) differed significantly in response magnitude, with 159 genes showing a stronger response in COVID-19 and 189 in TB. COVID-19 samples showed reduced expression of the maturation markers CD10 (*MME*) and CD16 (*FCGR3A*), alongside elevated expression of granule-cargo and NET-associated genes, including *MPO* and *ELANE*, consistent with a greater contribution from immature, emergency granulopoiesis-derived neutrophil populations. In contrast, TB samples showed higher maturation-associated, receptor, oxidative, and recruitment responses, consistent with a relatively mature receptor-competent neutrophil transcriptional signature. Independent COVID-19 Ward and pulmonary TB cohorts showed directional agreement for maturation and NET-associated signatures (100% and 90% concordance, respectively), while chemotaxis and select receptor and degranulation genes showed weaker reproducibility indicating that maturation represents the most consistently reproduced axis of distinction between the two diseases. Together, these findings identify maturation-associated transcription as the most consistently reproduced axis distinguishing COVID-19 from TB. These signatures provide candidate biomarkers for future studies linking neutrophil states to clinical outcomes and informing host-directed therapeutic strategies that preserve antimicrobial function while limiting immunopathology.

## 1 Introduction

Neutrophils are abundant, short-lived innate immune effector cells that comprise 40–70% of circulating leukocytes and can promote antimicrobial defense or contribute to inflammatory pathology. While traditionally viewed as a homogeneous population of immune cells, recent studies have revealed substantial heterogeneity and plasticity within neutrophils (1). For example, like macrophages, neutrophils are now considered to undergo differential polarization into N1 pro-inflammatory and N2 anti-inflammatory neutrophils characterized by distinct cell surface markers and chemotactic behavior. Neutrophil heterogeneity also extends to their buoyancy. While low-density neutrophils (LDNs) show activated phenotype with enhanced production of reactive oxygen species (ROS), cytokines and neutrophil extracellular traps (NETs) and increased expression of adhesion molecules and chemokine receptors, normal-density neutrophils (NDNs) show more robust antimicrobial functions, phagocytosis, and degranulation. CD10 and CD16 are commonly used surface markers to distinguish mature from immature neutrophil populations. CD10 is acquired during the later stages of granulocytic differentiation, and CD10 positive neutrophils generally represent terminally differentiated mature cells (2). CD16 expression also increases with neutrophil maturation. Mature circulating neutrophils typically exhibit high CD16 expression while immature neutrophils show reduced or absent expression (3). These differences in neutrophil population highlight their distinct roles in disease pathophysiology and progression.

In bacterial infections, neutrophil responses have been associated with both early protection and disease severity depending on timing, tissue context, and functional state (4, 5). Protective associations are most consistently observed during the early phase of infection at barrier tissues, particularly in cytokine environments that promote rapid neutrophil recruitment and antimicrobial function (6, 7). These associations are not merely defined by neutrophil abundance alone but by coordinated differences in timing, localization, and effector capacity, supporting the concept that neutrophils represent an adaptable correlate of protection rather than a rigid marker of infection (8, 9).

Neutrophil responses play a central role in host defense against viral pathogens (10). However, in contrast to many bacterial infections, neutrophil-associated signatures in viral diseases more frequently correlate with disease severity rather than protection (11). Across multiple viral infections, excessive, sustained, or dysregulated neutrophil activation has been associated with tissue damage, vascular dysfunction, and poor clinical outcomes (12, 13). These observations suggest that, in viral contexts, neutrophils often serve as a negative correlate of protection, highlighting the importance of temporal and functional constraints on neutrophil programming (8). This dysregulated response aligns with the characterization of severe COVID-19 that shows immature neutrophil populations in peripheral blood (14). This phenotype is consistent with emergency granulopoiesis, a cytokine-driven release of immature neutrophils from the bone marrow during severe infection.

Against this background, COVID-19 and tuberculosis (TB) provide a direct comparison to test whether neutrophil maturation state distinguishes programs associated with inflammatory pathology and antimicrobial host defense. COVID-19 is characterized by acute systemic inflammation and emergency granulopoiesis, whereas tuberculosis reflects chronic granulomatous inflammation with persistent host-pathogen interactions (14, 15). In TB, neutrophils can contribute to antimicrobial host defense, but excessive accumulation during active disease is also associated with greater bacterial burden and tissue pathology (16, 17). Despite these differences, both diseases exhibit neutrophil-associated tissue pathology, altered oxidative responses, and dysregulated inflammatory signaling. These contrasting immunological contexts suggest that COVID-19 and TB recruit neutrophils at different stages of maturation, and that these maturation states consequently determine their distinct functional programs and pathological outcome. It remains unclear whether the major transcriptional differences between these diseases reflect distinct neutrophil maturation states or disease-specific functional reprogramming within similar maturation states. We therefore hypothesized that COVID-19 and TB produce distinct neutrophil maturation states that contribute to their differences in effector functions, with COVID-19 enriched for immature neutrophils and TB for relatively mature neutrophils. We predicted the differences in degranulation, receptor signaling, NETosis, and chemotaxis would align with established maturation markers including CD10 and CD16.

To address this question, we performed an integrated bulk RNA-Seq analysis of six independent COVID-19 and TB datasets using the standardized discovery workflow summarized in Figure 1. We evaluated shared responses and disease-associated differences across maturation, degranulation, receptors/activation, NET-associated genes, and chemotaxis. We then assessed the generalizability of these signatures in an independent pulmonary-TB cohort and a COVID-19 neutrophil dataset restricted to hospital Ward patients for validation.

**Figure 1.**
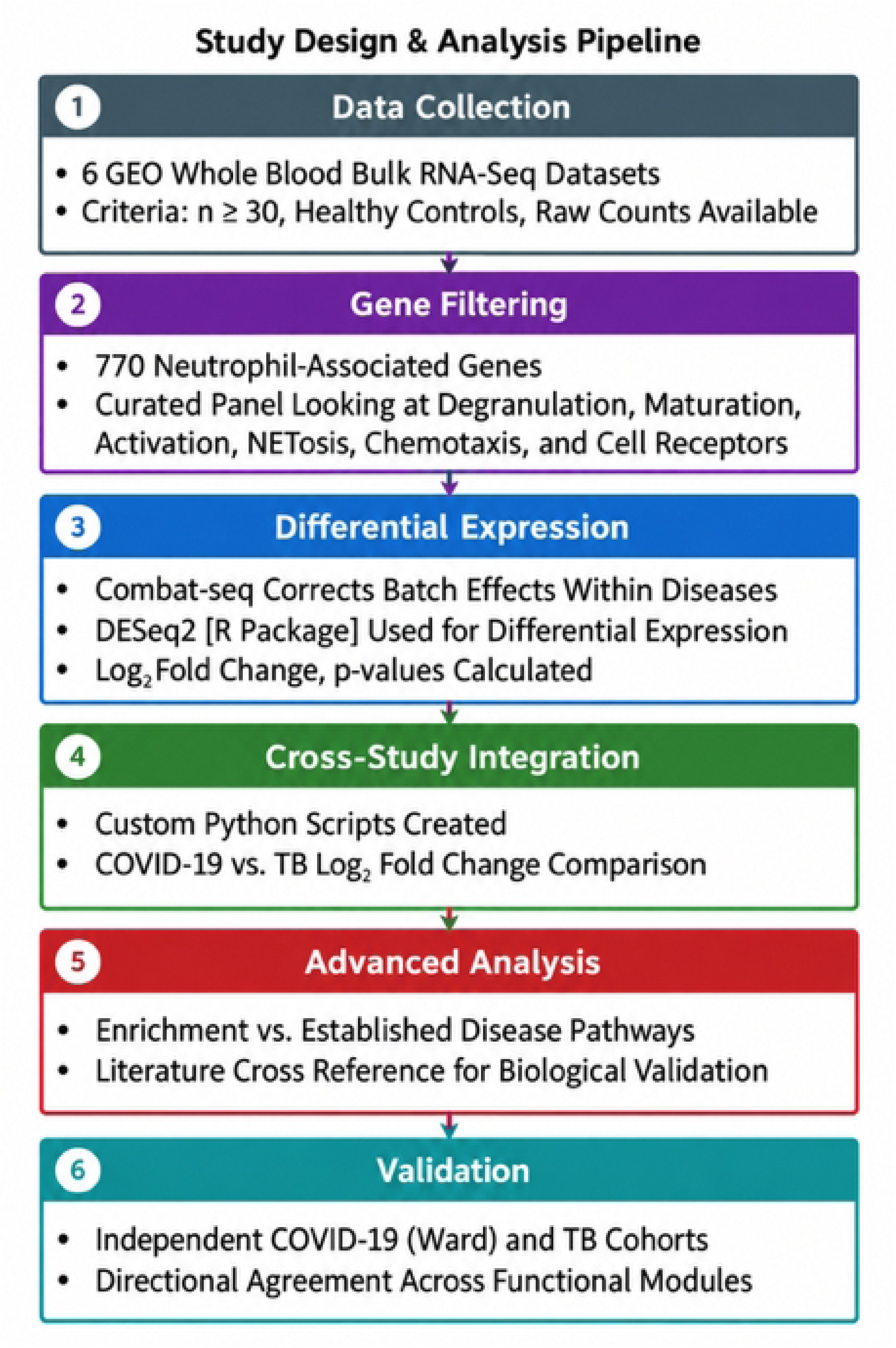
Schematic workflow of the integrative transcriptomic study. Six publicly available whole-blood RNA sequencing studies, comprising three COVID-19 and three active tuberculosis (TB) cohorts, were selected from the GEO database according to predefined criteria (Figure S1). A 770-gene panel targeting neutrophil-associated pathways was curated from GSEA, Reactome, KEGG databases, literature-derived gene sets, and additional manually curated genes. The analysis workflow included batch effect correction (ComBat-Seq), differential gene expression analysis (DESeq2), and cross-disease integration to identify shared and disease-differential neutrophil transcriptional signatures.

## 2 Materials and Methods

### 2.1 GEO dataset selection

Bulk RNA sequencing datasets were obtained from the Gene Expression Omnibus (GEO) repository. Searches to identify discovery and independent validation datasets were conducted between March 30, 2026, and October 7, 2026 using combinations of disease terms (“COVID-19” or “SARS-CoV-2”; “tuberculosis” or “Mycobacterium tuberculosis”), sequencing terms (“RNA-seq” or “bulk RNA-seq”), tissue terms (“blood”, “whole blood", or "human-samples"), and neutrophil-related terms (“neutrophil,” “granulocyte,” “PMN,” “CD15,” “low-density,” or “LDN”).

Datasets were eligible if they included at least 30 total patients, whole-blood bulk RNA sequencing data, raw gene expression data, and an appropriate comparator group: uninfected healthy controls for COVID-19 and latent tuberculosis infection (LTBI) controls for active TB. Control cohorts were evaluated against the NeutGX database (18). FPKM values were compared individually, gene by gene, with the corresponding NeutGX reference values (18). Comparisons were qualitative, without a predefined numerical exclusion threshold. Formal statistical testing was not performed because reference variability could not be characterized from the available NeutGX data. The selection workflow is shown in Figure 1.

The discovery list contained 19 accession records. After removing a duplicate dataset, 18 records were reviewed and 12 excluded under the final eligibility criteria, leaving six discovery cohorts (Figure S1A). Eleven datasets were excluded because their data types or sampled tissues did not meet the inclusion criteria, and one was excluded because its control samples were inconsistent with the NeutGX reference database. The six included datasets were as follows: three COVID-19 cohorts, GSE163151 (19), GSE217948 (20), and GSE161731 (21), and three active-TB cohorts, GSE107991 and GSE107992 (22) and GSE101705 (23). Only whole-blood samples were included. For GSE161731, only the first available sample from each participant was retained to avoid repeated measures during the period of rapidly changing neutrophil activity. Active TB was compared with LTBI in the discovery cohorts, as large whole-blood TB datasets frequently lack uninfected healthy comparator groups. Dataset characteristics, eligibility decisions, and final analytic sample counts are summarized in Table S1.

For independent validation, eleven candidate sources were reviewed and nine did not meet our selection criteria, leaving two validation cohorts (Figure S1B). Nine studies were excluded because of incompatible data, small sample sizes, or because of therapeutic intervention in the disease group. Independent validation used the COVID-19 dataset GSE234207 (24) and the pulmonary TB cohort reported by Hai et al. (2024). The discovery datasets were accessed for research purposes between March 30, 2026 and June 15, 2026, and the independent validation datasets were accessed between September 24, 2026 and Oct, 7 2026. All participant data were publicly available and de-identified before access. The authors had no access to information that could identify individual participants.

### 2.2 Neutrophil Gene Panel

A panel of neutrophil-associated genes was curated to represent neutrophil activation, neutrophil-mediated immunity, chemotaxis, degranulation, and NET-associated processes. Gene lists associated with neutrophil-mediated immunity, neutrophil activation, and chemotaxis were obtained using Gene Set Enrichment Analysis (GSEA) resources and comprised 21, 42, 85 genes, respectively. The GSEA neutrophil-associated genes were curated from the MSigDB Human Gene Ontology Biological Process collection (C5:GO), accessed June 17, 2026. The Reactome database contained 478 genes associated with degranulation. A NET-associated gene list was obtained from the KEGG database and had 193 genes. An additional panel of 69 genes from (25) was also included to provide additional coverage on NET formation. After literature survey, eight neutrophil-associated genes (*ANXA1*, *CEBPE*, *GFI1*, *MX1*, *NLRP3*, *STAT1*, *STAT3*, *STX4*), which were not part of the above gene sets, were further added. The various sets were then condensed to remove duplicates yielding 770 unique genes utilized for analysis. For analysis and reporting, the panel was organized into five functional modules: maturation, degranulation, receptors/activation, NET-associated genes, and chemotaxis. Module curation began with the broad gene panel assembled from the source datasets described above. Genes meeting the statistical significance criteria of our analysis pipeline were then assigned to modules according to their biological roles, as described in the original source datasets and additional neutrophil literature. Genes with documented roles in multiple processes were included in more than one module. The panel enabled focused analysis of neutrophil-associated transcriptional programs across both diseases. Figure S2 shows the distribution of genes across the source resources, including overlap among multiple neutrophil functions. The complete 770-gene panel, source annotations, and module assignments are provided in Table S2.

### 2.3 Differential Expression Analysis

The filtered datasets were exported as structured Excel files and analyzed in R version 4.5.1 using a pipeline that incorporated ComBat-Seq (sva v3.58.0) (26) batch correction followed by differential expression analysis with DESeq2 version 1.50.2 (27). ComBat-Seq was applied separately within the COVID-19 and TB datasets, with study accession specified as the batch variable and disease status as the biological condition. This approach reduced study-level batch effects while preserving within-disease case–control differences. Correcting all six studies together could confound disease identity with batch and attenuate the between-disease signal of interest. The batch-corrected count matrices were subsequently analyzed with DESeq2, which normalized the count data and performed differential expression testing between healthy or LTBI control groups and disease cohorts.

### 2.4 Cross-Study Integration and Visualization

Cross-study analysis was performed using a custom R script designed to integrate and compare differential expression results across datasets. Within each disease, DESeq2 generated log₂ fold-change values, nominal p-values, and Benjamini-Hochberg multiple testing-adjusted p-values relative to the relevant control group. For direct COVID-19 versus TB comparisons, an individual sample response value was calculated for each gene as the sample’s log₂-transformed normalized count (DESeq2 size-factor-normalized, batch-corrected counts, with a pseudocount of 1) minus the mean log₂-transformed normalized count of the relevant control group (healthy controls for COVID-19; LTBI for TB). Response values were pooled across studies within each disease, and Welch’s t-test was used to compare mean response values between COVID-19 and TB, accounting for unequal sample sizes and variances. The resulting p-values were adjusted using the Benjamini-Hochberg procedure, and genes with an adjusted p-value < 0.05 were classified as significantly different between diseases. These outputs were used to generate volcano plots and comparative visualizations of within-disease and cross-disease expression patterns.

### 2.5 Independent validation cohorts and response construction

COVID-19 validation data were obtained from the NCBI Gene Expression Omnibus dataset GSE234207, which profiled isolated neutrophils from healthy participants and patients hospitalized in wards or intensive care (24). The comparison used the 12 Ward samples, and 10 healthy-control samples. The 10 ICU samples were excluded to better align the validation cohort with the clinical profile of the COVID-19 patients in the test dataset. TB validation used whole-blood RNA-sequencing data from Hai et al. (2024), comprising 295 pulmonary-TB participants and 30 healthy controls from Vietnam. Tuberculous-meningitis samples were excluded. The filtered TB workbook contained 404 genes represented on a supplied variance-stabilized expression scale.

COVID-19 sample responses were calculated as the log_2_-transformed DESeq2 size-factor normalized count minus the corresponding healthy control mean (27). Normalization factors were estimated using the Ward samples and the COVID-19 healthy controls. For TB validation, the supplied dataset contained expression values that had already been normalized and variance-stabilized using DESeq2 by the original investigators (28). The validation response values were therefore both on log₂-based expression scales, but COVID-19 data were processed de novo from raw counts, whereas TB data entered the analysis after source-study normalization and variance stabilization (28). Each pulmonary-TB response was calculated by subtracting the healthy-control mean for that gene from the patient’s supplied expression value. Pulmonary-TB samples were compared with healthy controls using limma linear models and Benjamini–Hochberg-adjusted p-values. Genes were flagged when the adjusted p-value was below 0.05. The validation datasets also differed in cellular source, with isolated neutrophils used for COVID-19 and whole blood used for TB. Validation therefore emphasized the direction of the COVID-19–TB contrast rather than direct comparison of response magnitude.

### 2.6 Directional agreement and module comparison summaries

For each gene, we subtracted the mean TB response from the mean COVID-19 response in both the discovery and validation datasets. Positive differences favored COVID-19, whereas negative differences favored TB. Directional agreement was evaluated for genes in the original five module panels that had genes in both validation datasets and had a discovery adjusted Welch’s t-test p-value below 0.05. Validation significance was not required for inclusion. Direction was retained when the same disease had the higher mean response in both datasets and reversed when the disease with the higher mean response changed between datasets. Missing genes were excluded from the analysis. Directional concordance between discovery and validation cohorts was assessed for each module using a two-sided exact binomial test against a null expectation of 50% agreement (p = 0.5). For a module with *n* testable genes, the smallest attainable two-sided p-value is 2 × 0.5ⁿ. For the five gene maturation module, this minimum is 0.0625, so results for this module are interpreted descriptively.

### 2.7 Ethics statement

This study was reviewed by the Los Alamos National Laboratory Human Subjects Research Review Board (HSRRB), which determined that it did not constitute human subjects research. No participants were recruited or new human specimens collected. Individual informed consent was not required for this secondary analysis of publicly available, de-identified data.

## 3 Results

### 3.1 Study-level variation was reduced before integrated analysis

Prior to ComBat-Seq correction and dataset integration, study-level volcano plots demonstrated substantial variation in the magnitude and direction of differential expression (Figure S3A). The largest COVID-19 cohort (GSE217948) produced the broadest signature, comprising 107 significant differentially expressed genes (DEGs) and predominantly upregulated expression, whereas the other COVID-19 cohorts contained 31 and 22 significant DEGs with more mixed profiles. The three TB cohorts contained 28, 45, and 70 significant DEGs, respectively, and generally showed greater upregulation relative to LTBI. The DEGs were more consistent across the TB datasets (Figure S3B, S3C) with nine genes that were significant in all three TB studies and 31 that occurred in at least two datasets, compared with only one gene shared across all three COVID-19 studies and 20 occurring in at least two. The largest COVID-19 study also contained 87 study-specific genes, demonstrating greater heterogeneity among the COVID-19 datasets. Consistent with these findings, unsupervised hierarchical clustering separated the COVID-19 and TB studies into distinct disease-associated branches before additional transformation, while the two related TB cohorts clustered together (Figure S3). These pre-integration analyses identified reproducible disease-level structure, but also substantial study-specific variation, supporting the need for batch effect correction before integrated gene expression analysis.

ComBat-Seq adjustment was applied separately within each disease group, as described in Methods Section 2.3. Principal component analysis (PCA) was used to assess changes in study-associated clustering following correction. Figure S3D shows that before ComBat-Seq correction, samples clustered strongly according to their study of origin. After ComBat-Seq, samples from different studies showed substantially greater overlap, indicating that study-specific technical variation had been reduced while the underlying biological structure was retained (Figure S3E). This reduced clustering supported the effectiveness of ComBat-Seq in improving comparability across the individual TB and COVID-19 datasets.

### 3.2 COVID-19 and TB share directional neutrophil responses but differ in magnitude across functional modules

Across the 770 genes evaluated at the individual-sample level, COVID-19 and TB typically exhibited a shared directional transcriptional response but disease-specific differences in response magnitude. Direct comparison identified 348 genes (45.2%) with significantly different responses between the diseases, including 159 with higher responses in COVID-19 and 189 with higher responses in TB. As shown in Figure 2A and 2B, significant genes were distributed across all four quadrants with 149 increased and 92 decreased in both diseases, while 107 exhibited opposing responses. Among the genes that differed significantly between COVID-19 and TB (Figure 2C), Reactome contained the largest set (n = 244), followed by GSEA (n = 59), KEGG (n = 56), and the Zhang et al. gene list (n = 35). These counts are non-exclusive because many significant genes were found in more than one source database. For subsequent analyses, the neutrophil-associated genes were examined within five predefined functional modules: maturation, degranulation, receptors/activation, NET-associated genes, and chemotaxis. Module-specific directional agreement in the independent cohorts is summarized in Table 1 and integrated with the discovery findings in the following subsections. P-values were calculated using two-sided binomial tests to assess whether the proportion of genes retaining direction differed from the 50% expected chance. These p-values describe directional agreement across genes. They do not test whether individual genes are differentially expressed.

**Figure 2.**
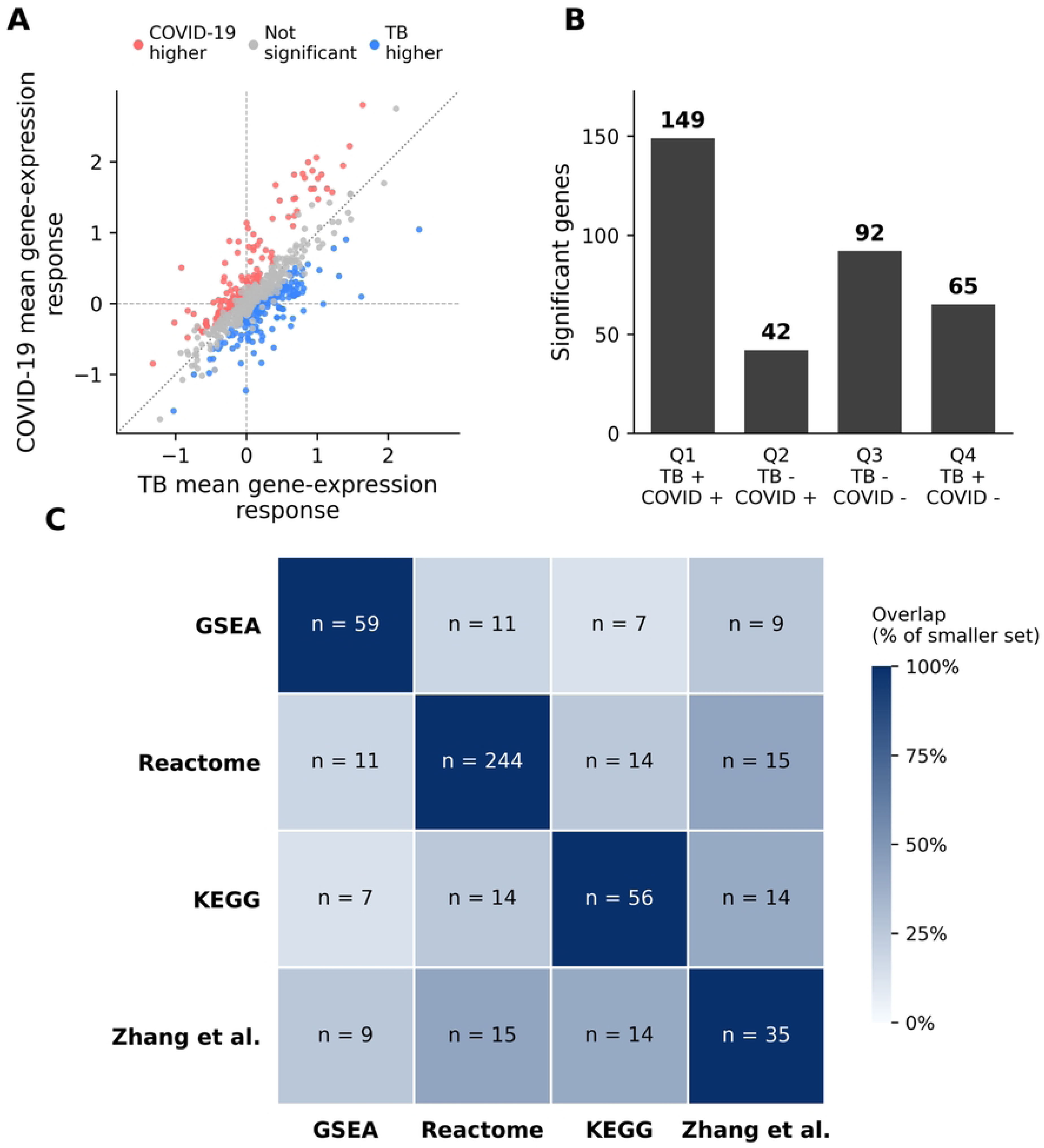
Cross-Disease comparison reveals maturation-associated transcriptomic divergence. **(A)** Scatter plot of log₂ fold change (log₂FC) for each gene with the x-axis representing the TB response and the y-axis representing the COVID-19 response. Red points indicate genes significantly upregulated in COVID-19 (adjusted p < 0.05); blue points indicate genes significantly upregulated in TB; grey points represent non-significant genes, **(B)** Bar chart showing the distribution of significant genes across the four quadrants from panel A, **(C)** Source distribution of significantly different genes showing overlap between genes across GSEA (neutrophil activation, immune response, and chemotaxis pathways), Reactome, and KEGG databases, and literature curation.

**Table 1.** Directional agreement across discovery and independent validation.

| **Module** | **Genes tested** | **Concordant** | **Reversed** | **p-value** | **Explanation** |
|---|---|---|---|---|---|
| Maturation | 5 | 5 (100%) | 0 (0%) | p = 0.0625<sup>a</sup> | All available markers retain direction; *CD47* and *PTPRC* unavailable. |
| Degranulation | 16 | 10 (62.5%) | 6 (37.5%) | p = 0.4545 | Retained cargo subset; reversals span primary/specific cargo, secretion, and *CYBB*. |
| Receptors and activation | 13 | 9 (69.2%) | 4 (30.8%) | p = 0.2668 | Fc-receptor contrasts retained; *CLEC6A*, *FPR1*, *FPR3*, and *CD44* reverse. |
| NET-associated | 10 | 9 (90%) | 1 (10%) | p = 0.0215 | Protease/cargo and recognition contrasts retained; *CYBB* reverses, *DEFA1* unavailable. |
| Chemotaxis | 9 | 4 (44.4%) | 5 (55.6%) | p = 1.000 | Selected recruitment contrasts reverse; cohort and processing influences remain possible. |
| Overall unique genes | 43 | 28 (65.1%) | 15 (34.9%) | p = 0.0660 | 65.1% concordance |

<sup>a</sup> p-values are two-sided exact binomial tests against 50% expected directional agreement. With five genes, the smallest attainable p-value is 2 × 0.5<sup>5</sup> = 0.0625, so the maturation module could not reach p < 0.05 even with 5/5 concordance. Genes missing from the validation datasets were excluded.

### 3.3 TB exhibits a stronger maturation-associated neutrophil signature than COVID-19

The neutrophil maturation branch showed a predominantly TB-weighted transcriptional pattern, with six of the seven genes showing stronger expression in TB than in COVID-19 as shown in Figure 3. *FCGR3A* (−0.08 (log₂ Fold Change) Cov, +0.52 TB, p-adj ≈ 0) and *MME* (−0.07 Cov, +0.50 TB, p-adj = 0.000025) demonstrated the largest between-disease separations, with mean response differences of approximately 0.59 and 0.57. *MME* encodes CD10, which distinguishes mature CD10⁺ from immature CD10⁻ neutrophil populations. Its selective increase in TB therefore supports a greater contribution from transcriptionally mature circulating neutrophils (2).

**Figure 3.**
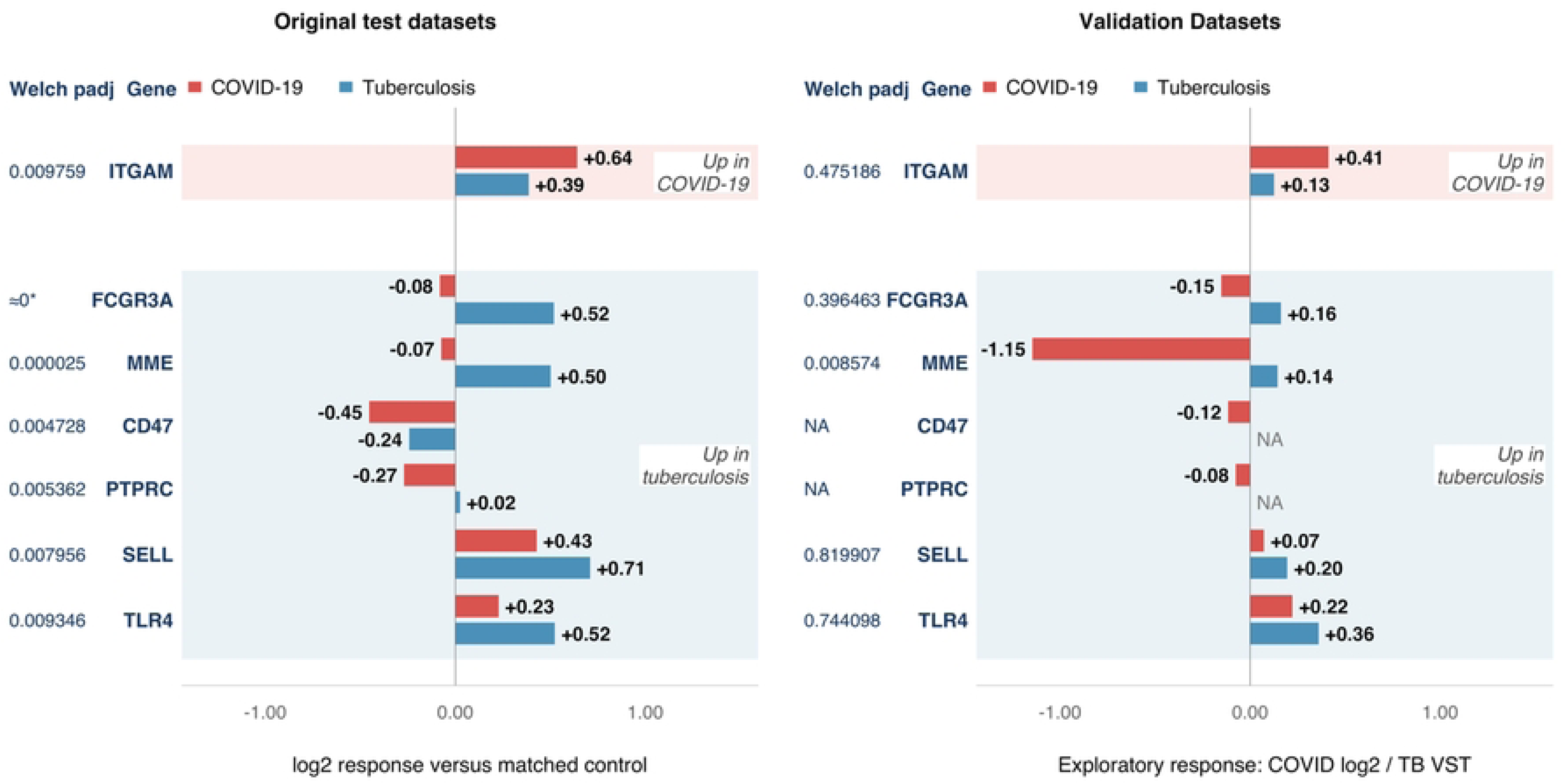
Neutrophil maturation marker expression reveals distinct maturation-associated transcriptional signatures in COVID-19 and tuberculosis. Differential expression of seven maturation-associated genes relative to disease-appropriate controls shows a predominantly TB-associated pattern. *FCGR3A*, *MME*, *SELL*, and *TLR4* are markedly upregulated in TB, while *PTPRC* remains near baseline and *CD47* is less strongly downregulated than in COVID-19. In contrast, COVID-19 shows reduced expression of *FCGR3A*, *MME*, *CD47*, and *PTPRC*, together with more modest increases in *SELL* and *TLR4*. *ITGAM* is the sole COVID-19-predominant marker, exhibiting greater upregulation in COVID-19 than in TB (+0.64 versus +0.39 log₂ fold change). These divergent expression patterns are consistent with disease-specific differences in neutrophil maturation. *Adjusted p-values displayed as 0 are shown as ≈0.

A similar TB-associated pattern was observed for *CD47* (−0.45 Cov, −0.24 TB, p-adj = 0.004728), *PTPRC* (−0.27 Cov, +0.03 TB, p-adj = 0.005362), *SELL* (+0.43 Cov, +0.71 TB, p-adj = 0.007956), and *TLR4* (+0.23 Cov, +0.52 TB, p-adj = 0.009346). CD47 is considered a supportive marker of neutrophil maturity because physiological expression facilitates the development of mature neutrophil subsets with functional chemotaxis, phagocytosis, reactive oxygen species production, and NET formation (29). *CD47* was suppressed in both diseases, but significantly less in TB. *PTPRC* encodes CD45, whose surface expression changes with human neutrophil development. CD45 expression accompanies progression toward mature neutrophils (30). *SELL* encodes L-selectin (CD62L), an adhesion receptor that mediates the initial tethering and rolling of neutrophils along the vascular endothelium. CD62L is rapidly shed following neutrophil activation and during transendothelial migration (31). Higher *TLR4* in TB may additionally indicate greater microbial-sensing and survival-signaling capacity, as TLR4 activation regulates neutrophil adhesion, respiratory burst, cytokine production, and life span (32). *ITGAM* (+0.64 Cov, +0.39 TB, p-adj = 0.009759) was the only COVID-19-dominant gene in this branch.

*ITGAM* produces CD11b, which combines with CD18 to form the Mac-1/CR3 integrin which supports neutrophil adhesion, migration, phagocytosis, and recognition of complement-opsonized targets (33). The greater *ITGAM* response in COVID-19 may reflect increased migratory activity as opposed to maturation. CD11b positive NET-forming neutrophil subsets have been associated with multiorgan dysfunction in severe COVID-19, although other flow cytometry studies found no overall increase in neutrophil CD11b relative to other infections or across COVID-19 severity groups (34, 35). The full data for this section is available in Table S3.

In the independent validation datasets, all five available maturation-associated genes retained their discovery direction (Figure 3). With only five maturation genes available for testing, the smallest attainable two-sided binomial p-value is 0.0625 (2 × 0.5⁵), so this module could not reach conventional significance (p < 0.05) even with complete concordance. The 5/5 agreement is therefore reported as a descriptive result (Table 1). *MME*, *FCGR3A*, *SELL*, and *TLR4* remained TB-weighted, whereas *ITGAM* remained COVID-19-weighted. The lower COVID-19 *MME* response remained consistent with a less-mature neutrophil contribution, whereas the *ITGAM* exception indicated that activation-related features did not follow a uniform maturation pattern. *CD47* and *PTPRC* were unavailable, so this result applies to five measured members of the seven-gene discovery panel.

Together, the discovery and independent-validation analyses identify the maturation-associated signature as the most consistently reproduced of the five modules, with complete directional concordance among the five testable genes. *MME, FCGR3A, SELL, TLR4,* and *ITGAM*, retained their discovery direction across the independent COVID-19 and TB cohorts (Table 1). Although *CD47* and *PTPRC* were unavailable in the TB validation dataset, complete directional retention among the remaining markers supports maturation-associated transcriptional state as a consistently observed axis distinguishing COVID-19 from TB. This finding does not establish that maturation alone determines downstream neutrophil function, but it identifies maturation-associated programs, rather than any individual effector pathway, as the most consistently reproduced difference between the diseases.

### 3.4 COVID-19 favors a broader degranulation-associated transcriptional program than TB

Neutrophil granule proteins are synthesized in stage-specific waves during granulopoiesis, thus degranulation-associated transcripts can reflect neutrophil maturation and subset composition in addition to cellular activation (36). Primary granule proteins are produced relatively early in development, whereas tertiary-granule proteins are synthesized during later maturation stages (36). The genes analyzed in this section are shown in Figure 4. Across the degranulation-associated genes identified by the individual-sample pipeline, both COVID-19 and TB exhibited a substantial neutrophil granule-associated transcriptional response relative to healthy controls, but the distribution of that response across functional modules differed markedly. Comparable degranulation-related transcriptional programs have been detected in both diseases, although such RNA-level signatures may reflect a combination of neutrophil activation, developmental state, subset composition, and changes in the abundance of circulating neutrophils (37). COVID-19 has specifically been associated with increased expression of neutrophil-degranulation genes (37). Active TB is characterized by a neutrophil-driven interferon inducible blood transcriptional signature, indicating that neutrophils contribute substantially to its gene expression profile (38).

**Figure 4.**
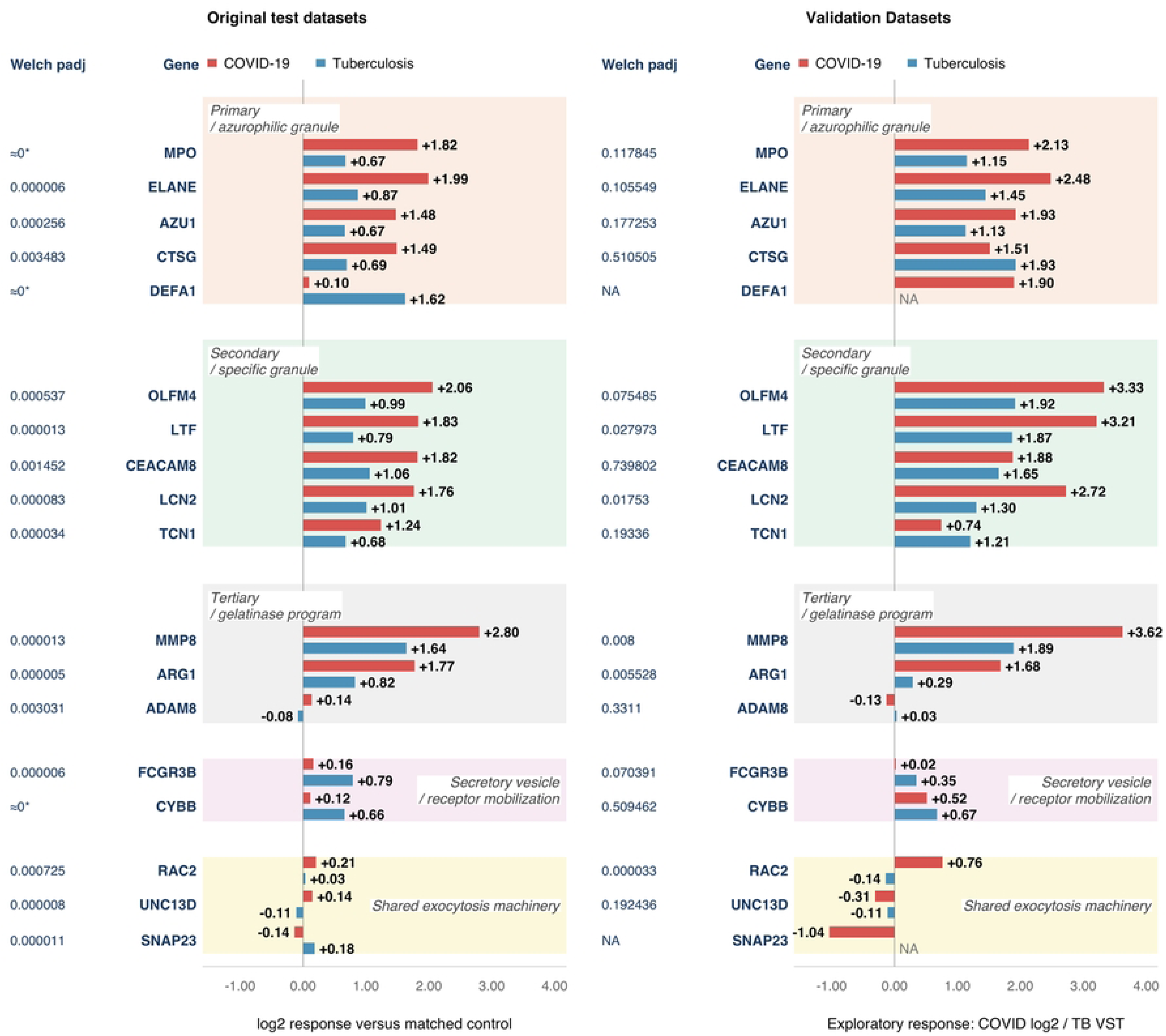
COVID-19 shows a broad granule-associated transcriptional program relative to TB. Comparative analysis of degranulation-associated gene expression organized by neutrophil granule subtype (primary, secondary, tertiary granules, secretory vesicles/receptor mobilization, and shared exocytosis machinery). COVID-19 samples demonstrate robust upregulation of primary, secondary, and tertiary granule genes, consistent with a broad granule-cargo transcriptional program. In contrast, TB samples show preferential upregulation of secretory vesicle components and the primary granule gene *DEFA1* (alpha-defensin 1), indicating a distinct antimicrobial response pattern. Genes encoding core exocytosis machinery show small magnitude but statistically significant differential expression, suggesting the disease distinction lies primarily in granule cargo rather than exocytosis machinery. *Adjusted p-values displayed as 0 are shown as ≈0.

Within the primary/azurophilic granule module, COVID-19 produced substantially larger mean responses for *MPO* (+1.82 Cov, +0.67 TB, p-adj ≈ 0), *ELANE* (+1.99 Cov, +0.87 TB, p-adj = 0.000006), *AZU1* (+1.48 Cov, +0.67 TB, p-adj = 0.000256), and *CTSG* (+1.49 Cov, +0.69 TB, p-adj = 0.003483), indicating a coordinated elevation of primary granules. These genes, together with *DEFA1*, encode classical azurophilic granule proteins whose expression is initiated during early granulopoiesis around the promyelocyte stage (36). In contrast, *DEFA1* (+0.10 Cov, +1.62 TB, p-adj ≈ 0) was strongly TB-dominant, demonstrating that the primary-granule pattern was not uniform and that TB preferentially emphasized an α-defensin component of the antimicrobial response.

The secondary/specific granule module provided the clearest and most consistent COVID-19-skewed pattern. *OLFM4* (+2.06 Cov, +0.99 TB, p-adj = 0.000537), *LTF* (+1.83 Cov, +0.79 TB, p-adj = 0.000013), *CEACAM8* (+1.82 Cov, +1.06 TB, p-adj = 0.001452), *LCN2* (+1.76 Cov, +1.01 TB, p-adj = 0.000083), and *TCN1* (+1.24 Cov, +0.68 TB, p-adj = 0.000034) were increased in both diseases but were consistently higher in COVID-19. These genes encode proteins associated with specific granule biogenesis or mobilization during the intermediate and late stages of neutrophil differentiation (36). This elevation is consistent with a stronger COVID-associated specific-granule transcriptional program involving the expression of antimicrobial nutrient-sequestering proteins, granule cargo, and membrane-associated activation factors (39). The tertiary granule cargo genes were also COVID-19 dominant, led by *MMP8* (+2.80 Cov, +1.64 TB, p-adj = 0.000013), which had the largest mean response of any gene in this set, whereas *ADAM8* (+0.14 Cov, −0.08 TB, p-adj = 0.003031) showed a statistically significant but comparatively modest difference.

In contrast to the granule-cargo modules, the secretory-vesicle/receptor-mobilization module favored TB. *FCGR3B* (+0.16 Cov, +0.79 TB, p-adj = 0.000006) and *CYBB* (+0.12 Cov, +0.66 TB, p-adj ≈ 0) were both substantially higher in TB, suggesting relatively greater emphasis on mobilizable Fc-receptor and NADPH-oxidase membrane components rather than broad granule-cargo transcription. Fc-receptor acquisition is linked to postmitotic neutrophil development and becomes most prominent with nuclear segmentation and terminal maturation, making the TB-dominant FCGR3B response consistent with the greater mature-neutrophil signal observed in the maturation branch (40).

Finally, the shared exocytosis machinery displayed lower-amplitude and directionally mixed changes. *RAC2* (+0.21 Cov, +0.03 TB, p-adj = 0.000725) and *UNC13D* (+0.14 Cov, −0.11 TB, p-adj = 0.000008) favored COVID-19, whereas *SNAP23* (−0.14 Cov, +0.18 TB, p-adj = 0.000011) favored TB. The small and opposing transcript changes observed here suggest that the principal distinction between the diseases lies in granule-cargo composition and neutrophil-state rather than exocytosis machinery. These findings support a broader and more intense degranulation-associated cargo program in COVID-19, particularly across secondary granules and matrix-remodeling components, whereas TB exhibits a more selective profile characterized by α-defensin expression and enhanced receptor/oxidase mobilization (41). When integrated with the maturation branch, the COVID-19 pattern showing strong expression of granule-biosynthesis genes together with lower *MME/CD10* and *FCGR3A* is compatible with emergency granulopoiesis and less-mature neutrophil states (14). The full data for this section are available in Table S4.

Independent validation separated the broad discovery degranulation pattern into retained and reversing components. Ten of the sixteen available genes retained direction, while six reversed (Figure 4). *ADAM8*, *UNC13D*, *CEACAM8*, *TCN1*, and *CTSG* were COVID-19 dominant in discovery but TB-dominant in validation. *CYBB* had an opposing reversal. *ADAM8* and *UNC13D* had small responses near zero, and modest differences between diseases in both comparisons. Given the relatively small COVID-19 validation cohort, these two reversals may reflect sampling variation rather than a substantial biological shift. The larger cargo changes provide a stronger basis for examining biological context than these two near-zero reversals.

The five reversing genes are functionally linked through neutrophil granule cargo, membrane mobilization, and secretion. *CTSG* encodes the primary/azurophilic granule protease cathepsin G (36), whereas the protein encoded by *TCN1* is a secondary/specific-granule marker released during neutrophil activation (42). CEACAM8, also known as CD66b, is predominantly stored in secondary granules (43). ADAM8 is a granule-associated disintegrin and metalloproteinase (44). Both proteins can move to the neutrophil surface upon activation. *UNC13D* encodes Munc13-4, which regulates granule secretion (45). These roles span several granule compartments, suggesting that their shared reversal may reflect differences in neutrophil developmental state or inflammatory priming rather than changes confined to one granule subtype.

*CTSG* showed a larger shift, combining a higher reported TB response with a lower COVID-19 response. The TB increases in *MPO*, *ELANE*, and *AZU1* place this change within a broader increase in primary-granule-associated responses, although those genes remained COVID-19-dominant. This pattern could reflect differences in the contribution of granule-producing neutrophil populations to whole-blood transcription (46). *CYBB*, in contrast, encodes NOX2, the catalytic component of the phagocyte NADPH oxidase complex involved in reactive oxygen species production. Its reversal primarily reflected a higher COVID-19 response, while the reported TB response remained similar. Increased *CYBB* transcription has also been reported in circulating low-density neutrophils from patients with severe COVID-19, providing a possible link to differences in inflammatory cell states of hospitalized patients (47). Thus, *CTSG* and *CYBB* may reflect changes in granule-cargo and oxidative machinery.

Pulmonary inflammatory burden provides a possible context for the TB-weighted cargo responses. In the validation source cohort, 129 of 270 participants with available chest radiographs had pulmonary cavities (28). Independent whole-blood analyses have linked greater PET-CT-defined lung damage to stronger degranulation- and NET-associated transcriptional signatures, including additional granule-cargo genes (48). Human pulmonary-TB studies also show neutrophils around cavity walls and enhanced *MMP8* secretion associated with radiological disease severity (49). Although *MMP8* retained the direction of the COVID-19–TB contrast in validation, it showed higher expression in the validation cohort than in the discovery cohort. These observations support a model where pulmonary inflammation alters both the abundance and the developmental or activation states of circulating neutrophils. A stronger contribution from granule-producing or inflammatory myeloid populations could consequently increase selected TB cargo responses and change their relative weighting against COVID-19.

Higher reported TB responses also extended across the five secondary/specific-granule-associated genes, including *OLFM4*, *LTF*, and *LCN2*, as well as *CEACAM8* and *TCN1*. Although *OLFM4*, *LTF*, and *LCN2* remained COVID-19-dominant, their higher TB responses narrowed the differences between diseases, while *CEACAM8* and *TCN1* became TB-dominant. This broader pattern is compatible with a greater contribution from secondary-granule-associated transcription in the TB validation cohort, potentially reflecting inflammatory burden or differences in circulating neutrophil populations. Although the discovery datasets also included pulmonary TB, clinical severity was not matched across cohorts and cannot be assumed to have been comparable. In the validation cohort, nearly half of the patients with available chest radiographs had pulmonary cavities, indicating substantial destructive lung disease in a large subgroup (28). This clinical context makes differences in inflammatory burden a plausible contributor to the higher reported TB granule-associated responses. Differences in disease burden and circulating neutrophil composition may therefore help explain the observed transcriptional pattern, although comparator and technical differences prevent its direct attribution to severity or increased granule release.

Together, the discovery and validation analyses indicate that the degranulation-associated program is partly reproducible but should not be treated as a uniform disease-defining signature. Ten of sixteen available genes retained direction across cohorts, including *MPO, ELANE, AZU1, OLFM4, LTF, LCN2, MMP8, ARG1, RAC2,* and *FCGR3B* (Table 1). In contrast, *ADAM8, UNC13D, CEACAM8, TCN1,* and *CTSG*, reversed from COVID-19-weighted in discovery to TB-weighted in validation. These results identify a candidate stable core of granule-cargo and related transcriptional differences, alongside a context-sensitive subset of granule- and secretion-associated genes that may be influenced by inflammatory burden, disease stage, or circulating cell composition.

### 3.5 Receptor- and activation-associated transcription is enriched in TB relative to COVID-19

Neutrophil maturation establishes the transcriptional and functional machinery required for migration and antimicrobial effector activity, while inflammatory priming can remodel receptor surface expression and function (50). This branch is best interpreted as an activation program paired with maturation state rather than as a direct readout of maturation alone (50). The activation and cell surface receptor branch showed a predominantly TB-weighted transcriptional pattern, as shown in Figure 5.

**Figure 5.**
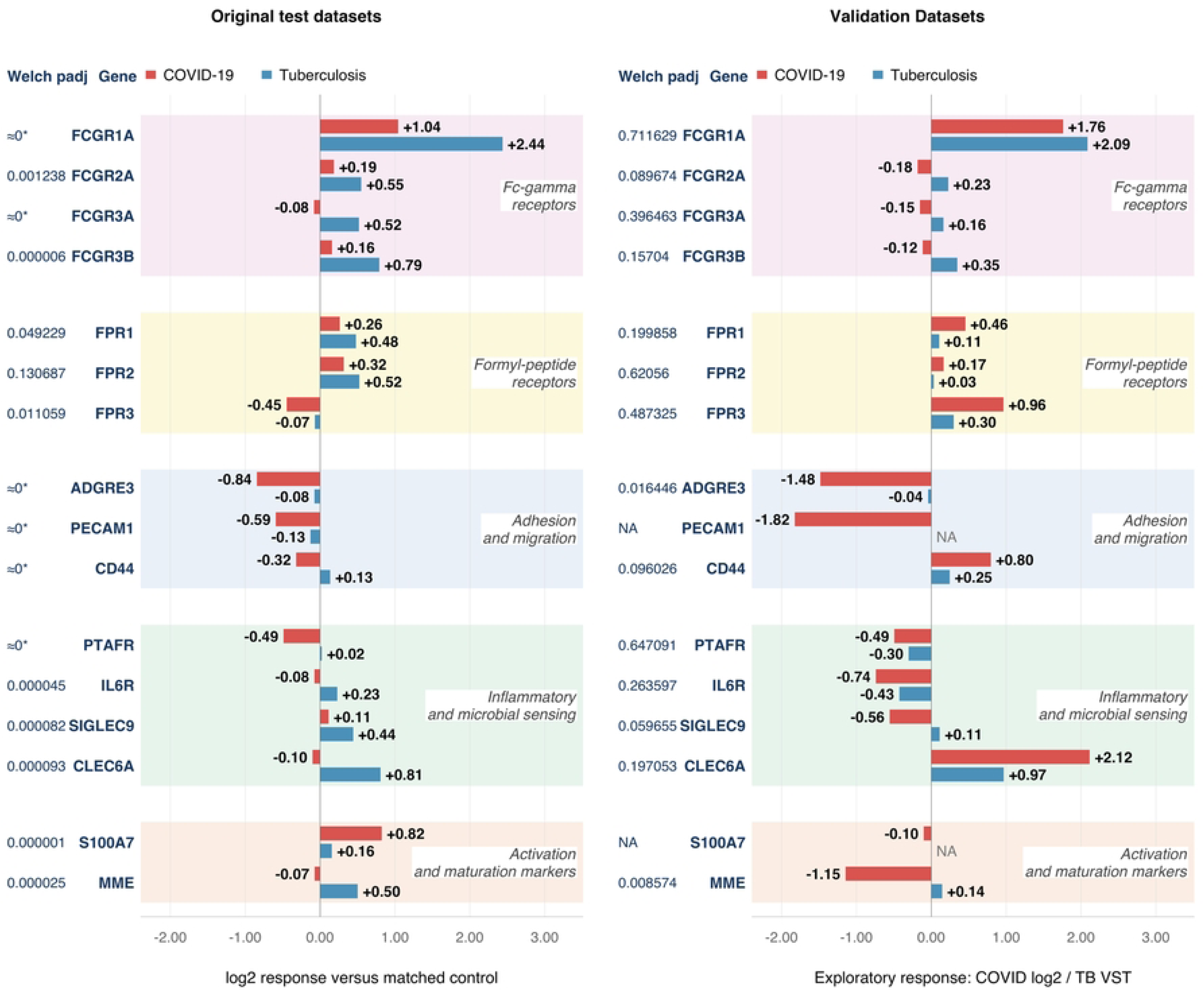
Neutrophil cell surface receptor and activation marker genes demonstrate a predominantly TB-weighted transcriptional signature. Comparative analysis of genes encoding cell surface proteins involved in neutrophil recognition, adhesion, and activation, organized by functional categories: Fcγ receptors, formyl peptide receptors, adhesion and migration markers, inflammatory and microbial pattern recognition receptors, and neutrophil activation markers. Fcγ receptors and formyl peptide receptors show higher responses in TB than in COVID-19, consistent with greater transcriptional emphasis on immune-complex recognition and formyl-peptide sensing.

All four Fcγ receptor genes had higher mean responses in TB than in COVID-19: *FCGR1A* (+1.04 Cov, +2.44 TB, p-adj ≈ 0), *FCGR2A* (+0.19 Cov, +0.55 TB, p-adj = 0.0012), *FCGR3A* (−0.08 Cov, +0.52 TB, p-adj ≈ 0), and *FCGR3B* (+0.16 Cov, +0.79 TB, p-adj = 0.000006). *FCGR1A* demonstrated the largest between-disease separation in this branch, with a mean-response difference of approximately 1.40. Fcγ receptor availability changes during postmitotic granulocytic development, supporting a developmental contribution to the receptor repertoire of circulating neutrophils. The large TB-associated increase in *FCGR1A*, encoding CD64, is more consistent with inflammatory activation than with terminal maturity alone (51). Interferon-γ can directly induce FCGR1A transcription in human neutrophils, while G-CSF can promote the appearance of CD64-positive mature granulocytes by acting on their precursors (52). FCGR2A and the neutrophil-enriched receptor FCGR3B participate in immune-complex recognition and can initiate calcium signaling, phagocytosis, oxidative burst, and degranulation (53). The coordinated TB-weighted pattern supports stronger Fc receptor–associated activation potential. Transcript abundance should not be equated directly with surface-receptor activity because neutrophil stimulation can mobilize receptors from intracellular stores or induce CD16 shedding independently of transcription (54). In addition, *FCGR1A*, *FCGR2A*, and particularly *FCGR3A* are expressed by other leukocyte populations, so their bulk-RNA signals may partly reflect differences in monocyte or natural–killer (NK) cell abundance rather than neutrophils alone (55).

A similar trend was observed among the formyl-peptide receptors, with *FPR1* (+0.26 Cov, +0.48 TB, p-adj = 0.049), *FPR2* (+0.32 Cov, +0.52 TB, p-adj = 0.13), and *FPR3* (−0.45 Cov, −0.07 TB, p-adj = 0.011) each having a higher response in TB, although the *FPR2* difference did not meet the adjusted significance threshold. Classical formyl-peptide receptor expression is acquired during normal human myeloid differentiation, supporting a developmental contribution to formyl-peptide sensing in mature granulocytes (56). The lower FPR1 and FPR2 responses in COVID-19 may indicate reduced transcriptional emphasis on formyl-peptide sensing, but they do not demonstrate a definitive loss of receptor function. FPR3 is more strongly associated with mononuclear myeloid populations than with circulating neutrophils, further emphasizing the contribution of other cellular groups to bulk expression (56).

Adhesion and migration-associated genes and inflammatory/microbial sensing receptors demonstrate preferential TB responses, supporting a broader receptor- and recognition-associated signature. Neutrophil activation markers show a divergent pattern, with *S100A7* showing COVID-19-predominant expression, while *MME*/*CD10* shows elevated expression in TB, suggesting a relatively mature neutrophil transcriptional signature. *Adjusted p-values displayed as 0 are shown as ≈0.

This broader TB-associated trend extended to the myeloid activation and migration genes *ADGRE3* (−0.84 Cov, −0.08 TB, p-adj ≈ 0), *PECAM1* (−0.59 Cov, −0.13 TB, p-adj ≈ 0), and *CD44* (−0.32 Cov, +0.13 TB, p-adj ≈ 0), as well as the inflammatory and microbial-sensing genes *PTAFR* (−0.49 Cov, +0.02 TB, p-adj ≈ 0), *IL6R* (−0.08 Cov, +0.23 TB, p-adj = 0.000045), *SIGLEC9* (+0.11 Cov, +0.44 TB, p-adj = 0.000082), and *CLEC6A* (−0.10 Cov, +0.81 TB, p-adj = 0.000093). *PECAM1* contributes to neutrophil trans-endothelial migration, whereas *SIGLEC9* functions as an inhibitory receptor capable of restraining neutrophil inflammatory responses (57, 58). Several genes in this group are expressed by monocytes, dendritic cells, endothelial cells, or other leukocytes. Their TB-weighted expression therefore supports a broader receptor-rich and migratory myeloid environment but cannot be attributed exclusively to neutrophil activation or maturation.

*MME* (−0.07 Cov, +0.50 TB, p-adj = 0.000025), encoding the mature-neutrophil marker CD10, was also higher in TB. CD10 distinguishes mature CD10-positive neutrophils from immature CD10-negative populations, providing the most direct maturation evidence within this branch (2). In the same study, mature CD10-positive neutrophils collected following G-CSF exposure exhibited an activated phenotype, showing that mature neutrophils can carry a strong activation program in an inflammatory setting (2).

*S100A7* (+0.82 Cov, +0.16 TB, p-adj = 0.000001) represented the only COVID-19 dominant gene in this cluster out of the 15 significant genes chosen for in-depth analysis. Although *S100A7* was more strongly induced in COVID-19, direct evidence linking this specific S100 gene to COVID-19 remains limited. *S100A8/A9* have been associated with severe disease, adverse clinical outcomes, and aggressive neutrophil activation (59, 60). Peripheral blood expression of *S100A4*, *S100A9*, and *S100A10* increases with COVID-19 severity (61).

The discovery receptor pattern complemented the maturation and degranulation findings: TB favored Fc-receptor and other recognition transcripts, while COVID-19 showed greater granule-cargo and protease responses. Neutrophil maturation helps determine which receptors cell express, while inflammation can alter receptor expression and trafficking can change receptor abundance on the cell surface. In whole-blood data, receptor-gene signals may also reflect contributions from other leukocytes. This distinction raised the question of whether the Fc-receptor component and broader sensing and adhesion responses would remain coupled in the independent comparison. The full discovery data for this section are available in Table S5.

The independent comparison retained nine of thirteen available discovery-significant receptor/activation contrasts (Figure 5). All four Fcγ receptor genes, *FCGR1A*, *FCGR2A*, *FCGR3A*, and *FCGR3B*, remained TB-weighted. In contrast, *CLEC6A*, *FPR1*, *FPR3*, and *CD44* shifted from TB-dominant in discovery to COVID-19-dominant in validation, occupying the upper-left quadrant of Figure S4. *CLEC6A* showed a particularly prominent reversal, whereas the discovery *FPR1* contrast was closer to zero. The independent evidence distinguished the retained Fc-receptor component from more variable pattern-recognition, chemotactic-sensing, and adhesion responses.

All four Fcγ receptor genes continued to show higher mean responses in TB than in COVID-19, but none of these differences were statistically significant after adjustment in validation (Figure 5). The sensing and adhesion genes still reversed direction after ICU patients were excluded, although differences in treatment, sampling time, severity, and cell composition may have contributed to these contrasts.

Together, the receptor and activation module showed moderate but incomplete reproducibility. Nine of thirteen available discovery-significant genes retained direction, including all four Fcγ receptor genes (*FCGR1A, FCGR2A, FCGR3A, FCGR3B*), which formed the clearest TB-weighted discovery (Table 1). However, the validation contrasts for these receptors did not remain statistically significant after adjustment, and *CLEC6A, FPR1, FPR3*, and *CD44* reversed direction. The reversal of pattern recognition, chemotactic sensing, and adhesion genes indicates that these components are more context-dependent than the directionally retained Fcγ-receptor pattern. Receptor-associated differences between COVID-19 and TB likely reflect both a relatively consistent receptor-related component and a variable layer shaped by activation, cellular composition, and inflammatory context. Surface flow cytometry will be needed to establish which receptor changes reflect neutrophil cell state versus shifts in circulating leukocyte populations.

### 3.6 COVID-19 and TB show divergent NET-associated protease/cargo and recognition/oxidative programs

In contrast to the granule-cargo transcriptional programs described above, NET formation is an effector response that cannot be attributed to maturation alone. Maturation may influence the availability of granule enzymes and oxidative machinery, but neutrophil abundance, subset composition, inflammation, pathogen recognition, and the extracellular environment are important determinants of the NET-associated transcriptional differences observed in Figure 6. Among the significant NETosis-associated genes identified by the analysis pipeline, COVID-19 and TB exhibited a modularly distinct pattern rather than uniform activation of a single NET-forming pathway. This distinction is important because NETs can be generated through multiple stimulus-dependent mechanisms, including both NOX2-dependent and NOX2-independent pathways (62).

**Figure 6.**
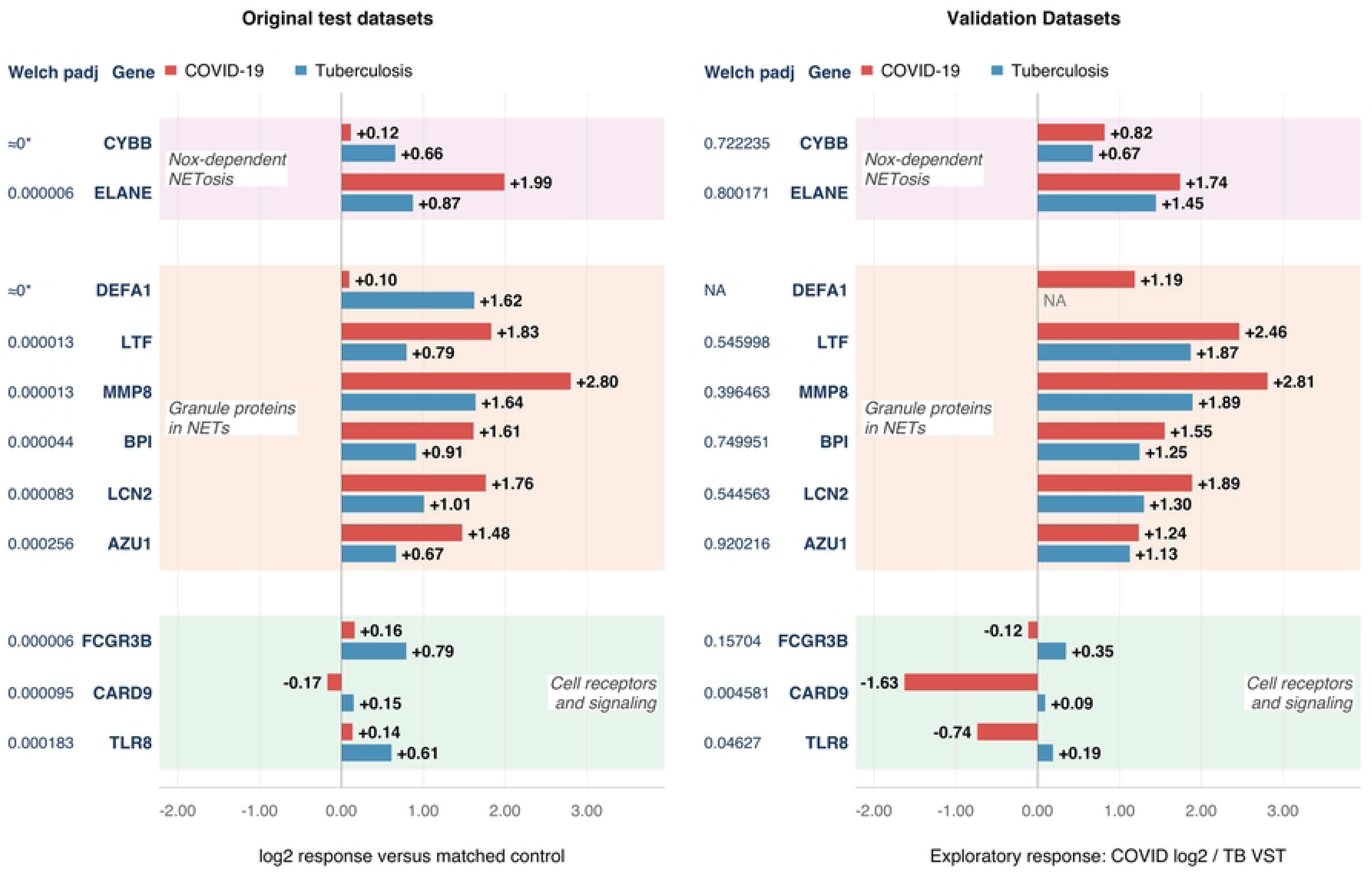
NET-associated gene expression reveals divergent protease/cargo and recognition/oxidative programs in COVID-19 versus TB. Comparative analysis of genes associated with NETosis and potential NET cargo, organized by functional categories: NOX2- and protease-associated pathway components, potential NET cargo, and receptor/recognition signaling genes. *CYBB* exhibits stronger upregulation in TB, while *ELANE* shows greater upregulation in COVID-19, indicating divergent oxidative and protease-associated transcriptional responses. Genes encoding potential NET cargo (*LTF*, *MMP8, BPI, LCN2, AZU1*) are predominantly upregulated in COVID-19, except for *DEFA1* which shows a stronger response in TB. Receptor and signaling genes associated with immune recognition and NET priming (*FCGR3B*, *CARD9*, *TLR8*) demonstrate preferential upregulation in TB, indicating a distinct upstream recognition and signaling signature. *Adjusted p-values displayed as 0 are shown as ≈0.

Within the NOX-dependent NETosis module, *ELANE* (+1.99 Cov, +0.87 TB, p-adj = 0.000006) was substantially more elevated in COVID-19, whereas *CYBB* (+0.12 Cov, +0.66 TB, p-adj ≈ 0) was more strongly elevated in TB. This opposing pattern is mechanistically informative because neutrophil elastase promotes histone degradation and chromatin degradation during NET formation, whereas the NOX2 catalytic component encoded by *CYBB* supports the reactive oxygen species production required for some forms of NETosis. Thus, COVID-19 preferentially expressed the granule-protease component of this pathway, while TB showed a comparatively stronger oxidative-enzyme signature (63). Neither disease displayed coordinated transcriptional elevation of the complete canonical NOX2-elastase pathway.

The clearest COVID-19-dominant pattern occurred within the granule-proteins associated with NETs module. *MMP8* (+2.80 Cov, +1.64 TB, p-adj = 0.000013) demonstrated the largest mean response in the entire NETosis-associated gene set, followed by *LTF* (+1.83 Cov, +0.79 TB, p-adj = 0.000013), *LCN2* (+1.76 Cov, +1.01 TB, p-adj = 0.000083), *BPI* (+1.61 Cov, +0.91 TB, p-adj = 0.000044), and *AZU1* (+1.48 Cov, +0.67 TB, p-adj = 0.000256), all of which were elevated in both diseases but consistently higher in COVID-19. The coordinated elevation of these genes indicates availability of potential NET cargo but does not establish that the encoded proteins were released on extracellular chromatin. This module also overlaps with the COVID-19-dominant degranulation program described in section 3.4. Much of the apparent COVID-19 NET signature may reflect broader induction of neutrophil granule cargo, increased neutrophil abundance, or altered developmental composition rather than selective activation of a dedicated NET-forming pathway. *DEFA1* (+0.10 Cov, +1.62 TB, p-adj ≈ 0) was a prominent exception, showing little mean elevation in COVID-19 but a strong response in TB.

The cell surface receptors and signaling module was uniformly more TB-dominant. *FCGR3B* (+0.16 Cov, +0.79 TB, p-adj = 0.000006), *TLR8* (+0.14 Cov, +0.61 TB, p-adj = 0.000183), and *CARD9* (−0.17 Cov, +0.15 TB, p-adj = 0.000095) each showed higher mean responses in TB. These genes are better interpreted as indicators of immune recognition and cellular priming than as direct components of the chromatin-release machinery. *FCGR3B* encodes CD16b, which contributes to neutrophil activation by immune complexes, while TLR8 detects microbial single-stranded RNA and CARD9 transduces signals from selected microbial-recognition receptors (64). This coordinated pattern suggests that the TB-associated NET signature may place greater transcriptional emphasis on immune recognition, receptor mobilization, and upstream signaling or priming, rather than on the broad granule-cargo induction observed in COVID-19.

*MPO* (+1.82 Cov, +0.67 TB, p-adj ≈ 0) was markedly higher in COVID-19. The encoded enzyme can cooperate with neutrophil elastase to facilitate chromatin modulation during select forms of NET formation (63). The COVID-19 elevation of *MPO* and *ELANE* provides stronger mechanistic support for enhanced protease-assisted chromatin decondensation than either gene alone, although both proteins also have NET-independent roles in degranulation and antimicrobial defense. The ROS-generation gene *HVCN1* (−0.64 Cov, −0.39 TB, p-adj = 0.000236), was reduced relative to controls in both diseases and was more strongly reduced in COVID-19.

The discovery findings combined COVID-19-weighted protease and granule-cargo responses with a more selective TB-weighted recognition, oxidase, and *DEFA1* pattern. The COVID-19 component is biologically consistent with experimental and clinical observations of circulating MPO-DNA complexes, spontaneous NET release, NET-containing pulmonary microthrombi, and induction of NETs by COVID-19 plasma (65). The independent comparison examined whether these cargo, recognition, and oxidative components retained their disease-associated direction. The full discovery data for this section are available in Table S6.

The NET-associated module showed the second-strongest level of reproducibility after maturation, with nine of ten available discovery-significant genes retaining direction (p = 0.0215; Table 1; Figure 6). The COVID-19-weighted protease and granule-cargo genes (*ELANE, LTF, MMP8, BPI, LCN2, AZU1*) and the TB-weighted receptor and recognition genes (*FCGR3B, CARD9, TLR8*) all held their original direction in the independent cohorts, indicating that the division between a COVID-19 protease/cargo signature and a TB receptor/recognition signature is directionally reproducible. *CYBB* was the sole reversal, shifting from TB-weighted in discovery to COVID-19-weighted in validation, and *DEFA1* could not be tested due to its absence from the validation TB dataset. The near-complete retention among NET-associated genes places this module alongside maturation as one of the two most stable distinguishing signals identified in this study, reinforcing that the protease/cargo and receptor/recognition transcriptional patterns carry reproducible disease-specific information.

### 3.7 TB favors a chemotaxis-associated recruitment and trafficking profile over COVID-19

Neutrophil chemotaxis reflects the interaction between developmental progress and inflammatory state. Neutrophil maturation establishes the receptor makeup and effector machinery, but receptor availability and activity are modified by inflammatory priming, chemoattractant signaling, endothelial adhesion, and transmigration (46, 50, 66). The following transcriptional pattern should not be interpreted as a direct measurement of either neutrophil maturity or migratory capacity alone, but rather a combination of these factors. The chemotaxis branch exhibited a highly consistent TB-weighted pattern, with 10 of the top 11 genes showing a greater mean response in TB than in COVID-19 (Figure 7).

**Figure 7.**
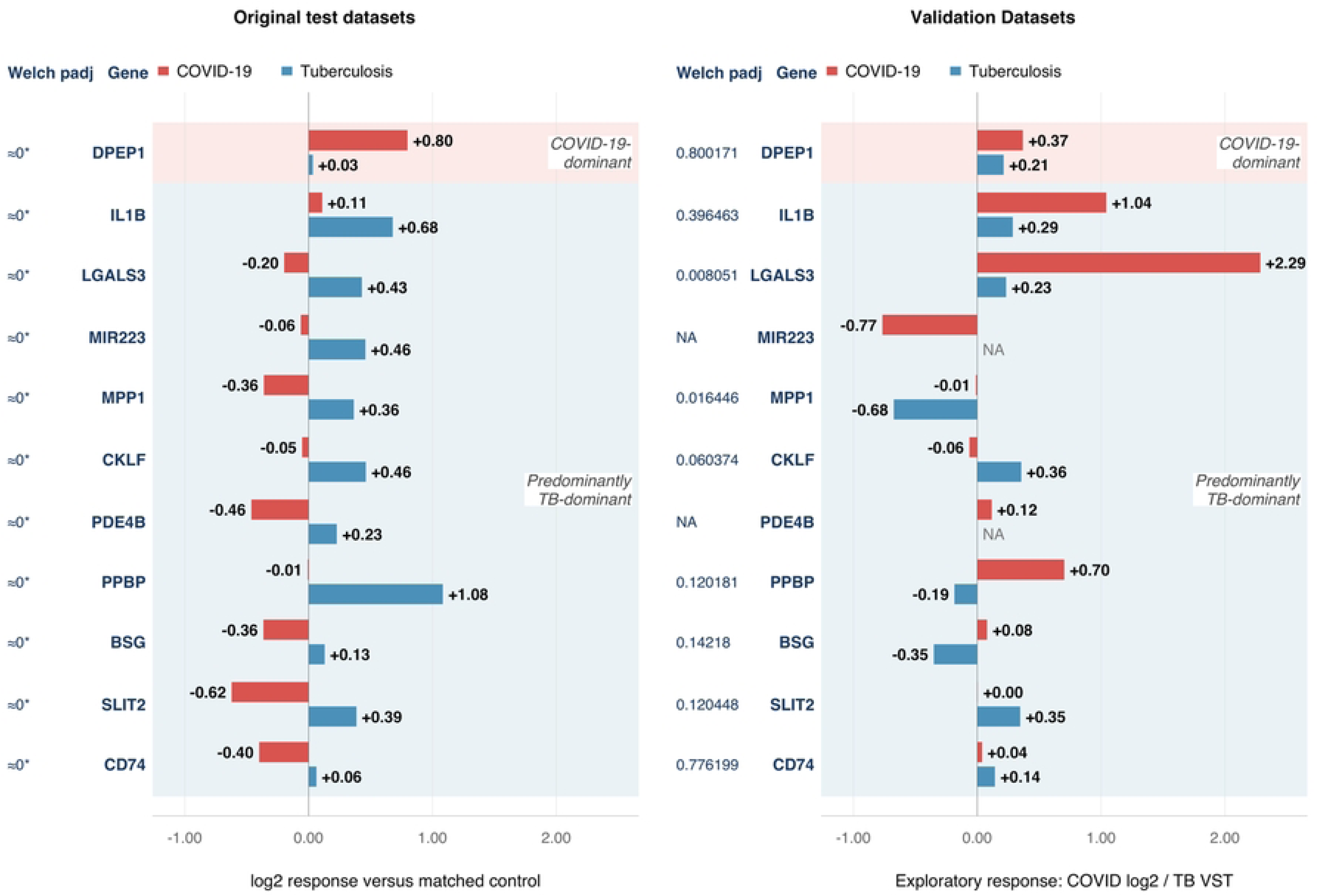
Chemotaxis-associated genes show a predominantly TB-weighted transcriptional pattern relative to COVID-19. Differential expression of the 11 discovery-significant chemotaxis-associated genes in COVID-19 and TB compared to disease-appropriate controls shows higher responses in TB for 10 of 11 genes (91%). In contrast, these same genes show predominantly lower responses in COVID-19, indicating a distinct recruitment and trafficking-associated transcriptional pattern. *DPEP1* represents the sole exception, showing COVID-19-predominant expression. *Adjusted p-values displayed as 0 are shown as ≈0.

The strongest TB-associated response was observed for *PPBP* (−0.01 Cov, +1.08 TB, p-adj ≈ 0), followed by *IL1B* (+0.11 Cov, +0.68 TB, p-adj ≈ 0), *CKLF* (−0.05 Cov, +0.46 TB, p-adj ≈ 0), *MIR223* (−0.06 Cov, +0.46 TB, p-adj ≈ 0), and *LGALS3* (−0.20 Cov, +0.43 TB, p-adj ≈ 0). *PPBP* encodes the platelet-derived chemokine CXCL7/NAP-2, which promotes neutrophil recruitment. IL-1β can promote recruitment indirectly by activating vascular and stromal cells, while CKLF1 has leukocyte-chemoattractant activity and galectin-3, encoded by *LGALS3*, facilitates neutrophil adhesion and extravasation (67, 68). Together, these genes suggest coordinated signaling among platelets, neutrophils, and the vascular compartment. *MIR223* provides the clearest maturation connection because miR-223 participates in a regulatory circuit controlling human granulocytic differentiation (69). Its higher expression in TB is compatible with the maturity-associated pattern observed in the maturation branch. *MIR223* alone cannot determine neutrophil maturity because it also regulates inflammatory activity.

A similar contrast was evident for *MPP1* (−0.36 Cov, +0.36 TB, p-adj ≈ 0), *PDE4B* (−0.46 Cov, +0.23 TB, p-adj ≈ 0), *BSG* (−0.36 Cov, +0.13 TB, p-adj ≈ 0), *SLIT2* (-0.62 Cov, +0.39 TB, p-adj ≈ 0), and *CD74* (−0.40 Cov, +0.06 TB, p-adj ≈ 0), each of which shifted from reduced expression in COVID-19 to a positive or approximately baseline response in TB. *PDE4B* regulates cAMP signaling and contributes to inflammatory neutrophil recruitment, although the supporting functional evidence is primarily preclinical (70). SLIT2 helps regulate neutrophil adhesion and migration through ROBO-receptor signaling (71). Different SLIT2 isoforms can either attract or repel neutrophils. Its higher expression in TB indicates altered regulation of neutrophil trafficking but does not establish whether migration was increased or inhibited (72). Direct neutrophil-chemotaxis evidence is limited for *MPP1*, *BSG*, and *CD74*.

*DPEP1* (+0.80 Cov, +0.03 TB, p-adj ≈ 0) was the only COVID-19-dominant gene in this branch. *DPEP1* can function as a neutrophil adhesion receptor in vascular and tissue compartments and thus may indicate a distinct adhesion pathway upregulated in COVID-19 (73). The TB-weighted profile puts greater transcriptional emphasis on platelet-mediated recruitment, inflammatory signaling, adhesion, and migration control. Maturation may contribute by providing a neutrophil population with greater receptor expression, consistent with the TB-weighted maturation and cell-receptor branches (46). This pattern complements the stronger receptor and oxidative signatures observed in TB. COVID-19 exhibited greater granule-cargo, protease, NET-associated, and *DPEP1*-mediated tissue-adhesion expression. The full data for this section are available in Table S7.

Independent validation preserved the COVID-19-weighted *DPEP1* contrast and the TB-weighted *CKLF*, *SLIT2*, and *CD74* contrasts (Figure 7). MIR223 and *PDE4B* were unavailable. Among the genes that reversed direction, *LGALS3* and *MPP1* met the adjusted validation significance threshold, while the other three did not. Several biological and design differences could contribute to the altered recruitment profile. Isolated-neutrophil COVID-19 samples and whole-blood TB samples capture different cellular contributions, and the TB comparator changed from LTBI in discovery to healthy controls in validation. Clinical isolates of *M. tuberculosis* can elicit different activation responses in human neutrophils (74), providing a biological basis for considering strain variation.

The chemotaxis module showed the weakest reproducibility of the five modules examined, with only four of nine available discovery-significant genes retaining direction. *PPBP, IL1B, MPP1,* and *BSG*, reversed from TB-weighted in discovery to COVID-19-weighted in validation, with *LGALS3* and *PPBP* showing particularly large shifts. This reversal pattern persisted even after ICU patients were excluded from the COVID-19 validation cohort, indicating that it was not explained solely by differences in disease severity. The reversals more likely reflect cohort-, sampling-, timing-, or strain-related differences in inflammatory and recruitment signaling that are harder to capture consistently across independent studies. The near-even split, and the particularly large reversals seen in *LGALS3* and *PPBP*, mark chemotaxis as the least reproducible of the five modules, indicating that chemotactic gene expression is governed more by contextual differences among cohorts than by a fixed disease identity.

## 4 Discussion

This study asked whether the contrasting neutrophil programs observed in COVID-19 and active TB can be explained by differences in maturation state rather than disease-specific functional reprogramming alone. The integrated analysis supports a partial developmental model: maturation-associated transcription was the most reproducible distinction, with COVID-19 showing an immature-associated program and TB a relatively mature, receptor-competent program. The NET-associated division was also strongly retained in independent cohorts, whereas degranulation, receptor/activation, and especially chemotaxis varied across cohorts. Therefore, maturation state appears to establish a reproducible starting point for divergent neutrophil programs, but activation and inflammatory context likely determine how that state is expressed and whether it contributes to host defense or pathology.

### 4.1 Maturation-associated neutrophil programs distinguish COVID-19 and TB

The main finding of this study is that neutrophil maturation represents an important point of divergence between COVID-19 and TB. COVID-19 exhibited a stronger immature-neutrophil signature, characterized by lower expression of mature-associated markers such as *MME* (CD10) and *FCGR3A*, together with consistent elevation of granule-cargo genes typically transcribed during neutrophil development (section 3.3). TB showed greater expression of maturation-associated markers and receptor-mediated recognition machinery, consistent with a more mature and receptor-competent circulating neutrophil transcriptional signature. The COVID-19 pattern is consistent with emergency granulopoiesis, a stress-induced response that causes inflammatory pressure to accelerate neutrophil production and releases developmentally immature cells into the circulation. This interpretation is supported by single-cell studies identifying circulating neutrophil precursors as evidence of emergency myelopoiesis in COVID-19 (14), developing neutrophil populations in the peripheral blood of patients (75), and hyperactivated immature neutrophil states associated with disease (76). The present analysis used bulk whole-blood RNA sequencing data, so these findings demonstrate a maturation-associated transcriptional pattern rather than directly proving that emergency granulopoiesis caused the observed differences.

This developmental model has explanatory power but only partial reach in that maturation state shapes the *starting* conditions of the neutrophil response, whereas activation, receptor availability, and tissue context can modify or override the expected developmental signature. Single-cell studies have identified circulating neutrophil precursors and immature neutrophil populations in COVID-19 (14). Phenotypic analyses have shown that these CD10⁻ populations can remain highly activated and exhibit increased secondary-granule release (76). The COVID-19 signature may therefore reflect both increased neutrophil production and a greater contribution from immature populations, whereas the higher *MME*/CD10 expression in TB suggests a relatively mature circulating compartment (2).

The NET-associated results indicate that maturation alone does not determine the direction of disease-specific transcriptional differences. COVID-19 favored protease and granule-cargo genes, while TB favored receptor, recognition, and oxidative genes (section 3.6). The opposing *ELANE* and *CYBB* responses indicate that neither disease uniformly increased every component of the canonical NOX2-dependent NET pathway. Experimental studies have demonstrated elevated NET formation and NET-containing pulmonary microthrombi in COVID-19, as well as NET formation in pulmonary TB (65, 77). Emergency granulopoiesis may increase the availability of granule enzymes that can later participate in NETs, but neutrophil abundance, inflammatory activation, platelet interactions, pathogen recognition, and the extracellular environment are more immediate determinants of NET release. Importantly, nine of ten available NET-associated genes retained their discovery direction in independent validation. This supports a reproducible division between a COVID-19-weighted protease/granule-cargo program and a TB-weighted receptor/recognition program, while the reversal of *CYBB* reinforces that neither profile represents uniform activation of a single NET-forming pathway.

The cell-receptor results further support a relatively mature and receptor-equipped neutrophil transcriptional signature in TB. The TB-weighted expression of Fcγ receptors and formyl-peptide receptors is consistent with a greater transcriptional emphasis on immune-complex recognition, microbial sensing, phagocytosis, and oxidative activation. Formyl-peptide receptor expression is acquired during myeloid differentiation, providing a direct developmental connection to neutrophil maturation (78). However, receptor expression is not determined by maturation alone. *FCGR1A*/CD64 can be induced by inflammatory and G-CSF-dependent signaling acting on myeloid precursors, indicating that the strong TB *FCGR1A* response likely also reflects activation (52). All four Fcγ-receptor genes retained their TB-weighted direction in independent validation, although none remained statistically significant after adjustment (section 3.5). In contrast, pattern-recognition, chemotactic-sensing, and adhesion genes showed less consistent directional retention. These findings support a directionally stable Fcγ-receptor component but indicate that the broader receptor signature is influenced by activation state and cohort context.

The degranulation findings provide a complementary connection to this developmental model. COVID-19 showed broad elevation across primary- and secondary-granule cargo and matrix-remodeling genes (section 3.4), reflecting the fact that granule components are produced in developmental waves during granulopoiesis, so their transcriptional abundance in whole blood may reflect the influx of cells retaining granule programming genes (79). Independent validation retained the direction of a subset of COVID-19-weighted cargo genes, but several cargo and secretion genes reversed direction. Thus, degranulation should not be interpreted as a single uniform disease-defining program. Instead, the results support a stable core granule-cargo signature alongside a context-sensitive subset.

Chemotaxis displayed a TB-weighted pattern spanning platelet-derived, inflammatory signaling, and adhesion genes (section 3.7), suggesting stronger coordination of platelet-derived recruitment, inflammatory signaling, adhesion, and extravasation. Maturation establishes the receptor machinery required for migration, but chemotactic behavior is modified by activation, receptor desensitization, endothelial contact, and prior trafficking (46, 66). The chemotaxis module showed the weakest independent reproducibility, with only four of nine discovery-significant genes retaining direction. It should therefore be interpreted as a context-dependent recruitment and trafficking signature rather than a stable maturation-associated distinction between the diseases.

### 4.2 Clinical implications

The neutrophil maturation-associated transcriptional signature associated with COVID-19 may also have important therapeutic implications. Dysregulated neutrophil responses have been implicated directly in COVID-19 acute respiratory distress syndrome (ARDS) (80). Excessive degranulation, NET formation, and neutrophil interactions can promote epithelial and endothelial injury. Veras et al. showed that COVID-19 induced NETs can directly promote pulmonary cell death (81). These findings suggest that maturation- and NET-associated markers could be evaluated as candidate biomarkers for stratifying neutrophil inflammatory states and monitoring future neutrophil-directed interventions. However, they do not establish that limiting immature neutrophil release or promoting maturation would mitigate COVID-19-associated ARDS. In TB, rather than an immature, protease-rich phenotype, active TB was characterized by a mature, receptor-competent neutrophil transcriptional signature, with elevated Fcγ and formyl-peptide receptor expression alongside enhanced oxidative-burst signaling (*CYBB*). The pattern suggests that future host-directed strategies for TB should account for receptor-mediated activation and oxidative capacity, rather than focusing on granulopoiesis alone. The same maturation markers that distinguished COVID-19 from TB, including *MME/CD10* and *FCGR3A*, may also hold candidate value as biomarkers for distinguishing active TB from latent infection, a distinction that remains clinically challenging with current diagnostics (82). These translational possibilities remain speculative and require functional studies and validation in well-matched active-TB, LTBI, and healthy-control cohorts.

### 4.3 Limitations

Several considerations should be taken into account when interpreting the findings of this study. The principal biological limitation is that whole-blood bulk RNA-seq cannot resolve changes in neutrophil subpopulation abundance from transcriptional changes within individual neutrophils. Neutrophils occupy a developmental spectrum and can enter circulation at different stages of maturity (83). Therefore, the observed signals may reflect different contributions from mature high-density neutrophils, immature low-density neutrophils, other myeloid populations, and non-neutrophil leukocytes. Patient-to-patient variation in the neutrophil-to-lymphocyte ratio (NLR) introduces additional uncertainty. NLR variation is associated with widespread differences in whole-blood expression that may not be fully captured by standard normalization approaches (84). Stronger neutrophil-associated signals may reflect greater neutrophil abundance or fewer lymphocytes rather than increased expression within individual neutrophils. Patient-level NLR data was not readily available for the included cohorts, preventing adjustment for this potential variation. Transcriptomic differences also cannot be assumed to correspond directly to protein abundance or neutrophil effector activity (85, 86). Although conducted in an LPS-stimulated lung epithelial model rather than neutrophils, Jacobsen et al. (2024) similarly demonstrated that transcriptional and protein-level immune responses may diverge (87).

The 770-gene panel used here is pre-selected and hypothesis-driven, and while it focuses the analysis on known neutrophil biology, it is necessarily incomplete and may miss novel or understudied effector programs. Validating the neutrophil maturation panel against single-cell RNA sequencing and flow cytometry data would help confirm that the transcriptional patterns observed reflect genuine cell-state shifts rather than dilution effects. Separately, lung tissue or bronchoalveolar transcriptomic data would be particularly informative for TB and COVID-19, as disease-specific signatures may be stronger and more apparent in respiratory tissues than in blood.

Combining six independent cohorts also introduced technical and clinical heterogeneity, including differences in geography, sequencing protocols, disease severity, cohort size, and sample-collection timing. ComBat-Seq reduced study-associated variation, but residual batch effects and population-specific immune background may remain. In addition, the timing of sampling relative to symptom onset was inconsistently documented, and differences in disease stage may contribute to variation in neutrophil abundance, maturation, and transcriptional activity. The discovery comparison also used different control populations: COVID-19 samples were compared with uninfected healthy controls, whereas active TB samples were compared with individuals with LTBI. Although these groups clustered together in Figure S5, LTBI may still be associated with immune or neutrophil priming, and control group differences could affect the magnitude or direction of individual disease-associated responses. Finally, the independent comparison used distinct sample types and control groups, so validation was interpreted primarily as directional agreement rather than exact replication of response magnitude. The retention of the maturation and NET-associated signatures across these independent cohorts supports the robustness of the central findings, whereas the weaker replication of other modules identifies findings that require further study.

### 4.4 Future directions

Future studies should focus on resolving the neutrophil population structure underlying the COVID-19 and TB transcriptional patterns identified in this analysis. Single-cell RNA sequencing could help distinguish changes in subpopulation proportions from transcriptional changes within those populations. The SCAHN atlas provides a reference for this work by defining immature granule-associated, mature, interferon-responsive, and inflammatory neutrophil programs (88). Comparing our panel with these atlas-derived signatures could guide marker selection for targeted single-cell and flow cytometry validation. Controlled studies using standardized sampling and well-characterized cohorts could further determine whether these signatures reflect differences in neutrophil maturation, activation, or functional state. Together, these approaches would clarify whether COVID-19 and TB are associated with distinct neutrophil population structures, disease-specific transcriptional reprogramming, or both.

## 5 Conclusions

This integrated analysis of six whole-blood RNA-sequencing cohorts demonstrates that COVID-19 and active TB, while sharing a broadly overlapping neutrophil transcriptional response, diverge substantially in maturation state and effector-program composition. While many of the 770 evaluated genes changed in the same direction, 348 of 770 evaluated genes (45.2%) differed significantly in response magnitude between the diseases with 159 stronger in COVID-19 and 189 stronger in TB. The most reproducible distinction was maturation-associated transcription: COVID-19 showed lower expression of *MME*/CD10 and *FCGR3A*/CD16 alongside broad granule-cargo and protease expression, whereas TB showed a relatively mature, receptor- and oxidative-associated signature. This pattern is consistent with a greater contribution from immature, emergency-granulopoiesis-derived neutrophil populations in COVID-19, but bulk whole-blood data cannot establish that emergency granulopoiesis caused these differences. Independent validation confirmed directional retention of the maturation and NET-associated signatures, whereas degranulation and receptor/activation signals were only partly reproducible and chemotaxis was strongly cohort dependent. Together, these findings identify maturation-associated transcriptional state as a reproducible but incomplete axis of divergence between COVID-19 and TB. Maturation state may establish the starting conditions for disease-specific neutrophil programs, whereas activation, receptor trafficking, and tissue context likely determine how those programs are expressed. These signatures provide candidate biomarkers for distinguishing neutrophil inflammatory states and prioritizing future host-directed strategies, pending validation by single-cell transcriptomics, flow cytometry, and functional assays in blood and respiratory tissue.

## 7 Author Contributions

SH: Writing – original draft, Data curation, Formal analysis, Methodology, Conceptualization. AS: Data curation, Methodology, Writing – review & editing, Supervision. JKS: Supervision, Conceptualization, Writing – review & editing, and Funding acquisition.

## 9 Acknowledgements

The authors thank Dr. Traci Pals and Dr. Bob Webb at DTRA for their support of this work. This work was performed at the U.S. Department of Energy’s Los Alamos National Laboratory. Los Alamos National Laboratory is operated by Triad National Security, LLC, for the National Nuclear Security Administration of U.S. Department of Energy (Contract No. 89233218CNA000001).

During preparation of this manuscript, the authors used ChatGPT (OpenAI; GPT-5.6 Sol and GPT-5.6 Tera) to assist with language editing and to draft R scripts used to generate Figures 2–7. The scripts were executed by the authors in R using the study’s DESeq2 and Welch’s t-test outputs. All plotted values were independently verified against the source differential-expression tables (Tables S3–S7) prior to inclusion. Generative AI was not used to generate or alter the underlying data or statistical analyses. All authors reviewed, revised, and verified the final manuscript and take full responsibility for its accuracy, originality, and integrity. The graphical abstract was created in BioRender. Harrington, S. (2026)

## 11 Supporting Information

**Figure S1. Study selection for discovery and independent validation.** (A) The discovery list contained 19 GEO accession records. Removal of one parent-record duplicate left 18 records for eligibility review. Twelve were outside the final criteria: 11 because of sample type or profiling method and one, GSE166190, following qualitative NeutGX comparison and review of family sampling. Six whole-blood bulk RNA-sequencing cohorts were retained. (B) Eleven candidate sources were reviewed for independent validation. Three were incompatible with the planned comparison and six others were not selected. The final validation cohorts comprised isolated neutrophils from 12 Ward COVID-19 patients and 10 healthy controls (GSE234207), and whole blood from 295 pulmonary TB patients and 30 healthy controls (Hai et al., 2024).

**Figure S2. Source distribution of the 770-gene panel used for differential expression analysis.** This depicts the overlap between genes curated from GSEA (including neutrophil activation, immune response, and chemotaxis pathways), Reactome, KEGG, and literature-based selections (Zhang et al., 2022). The diagram demonstrates both unique contributions from each database and shared genes across multiple annotation sources.

**Figure S3. Dataset quality control.** (A) Volcano plots showing differential gene expression for each of the six included datasets relative to their respective healthy control populations. Genes meeting the significance criteria are shown as upregulated or downregulated, while nonsignificant genes are shown in gray. (B–C) Venn diagrams illustrating overlap in significantly differentially expressed genes among the three COVID-19 datasets (B) and the three tuberculosis (TB) datasets (C), demonstrating both shared disease-associated transcriptional responses and study-specific signatures. (D–E) Principal component analysis (PCA) of all samples before (D) and after (E) cross-study normalization/batch correction. Prior to correction, samples show substantial clustering by individual study, whereas correction reduces study-associated separation while preserving broader disease-related transcriptional structure. Together, these analyses demonstrate consistent differential-expression signals across independent cohorts and support the integration of the six datasets for subsequent comparative analyses.

**Figure S4. Exploratory agreement of discovery and Ward-only validation contrasts.** The x-axis shows the discovery mean COVID-19-minus-TB response in log2 units; the y-axis shows the validation contrast calculated from COVID-19 log2 responses and TB variance-stabilized responses. Each point represents one of 43 unique discovery-significant genes available in both analyses. Circles indicate retained direction and triangles indicate reversed direction. Colors assign each gene to its first matching module for display; biological memberships overlap. Twenty-eight genes retained direction and fifteen reversed. The dashed y = x line is a graphical reference rather than evidence of calibrated effect-size equivalence. COVID-19 validation includes Ward patients only; ICU patients are excluded. TB disease and sequencing run are confounded, and the two disease responses use different transformed scales.

**Figure S5. Unsupervised hierarchical clustering demonstrates clear separation between COVID-19 and tuberculosis transcriptomic profiles.** Dendrogram showing hierarchical clustering of six datasets using the raw gene data prior to any data transformations. All COVID-19 studies (n=3) cluster together in one major branch, while all TB studies (n=3) cluster in a distinct branch, indicating robust disease-specific gene expression signatures. The clustering of LTBI with the healthy non-tuberculosis control shows that LTBI and healthy samples have similar profiles.

**Table S1. Characteristics of the six discovery cohorts.**

**Table S2. Complete gene list with source datasets.**

**Table S3. All significant maturation genes.** This table represents a list of all differentially expressed genes with a p-adj value < 0.05.

**Table S4. All significant degranulation genes.** This table represents a list of all differentially expressed genes with a p-adj value < 0.05.

**Table S5. All significant cell receptor genes.** This table represents a list of all differentially expressed genes with a p-adj value < 0.05.

**Table S6. All significant NETosis genes.** This table represents a list of all differentially expressed genes with a p-adj value < 0.05.

**Table S7. All significant chemotaxis genes.** This table represents a list of all differentially expressed genes with a p-adj value < 0.05.

## 12 Data Availability Statement

Publicly available datasets were analyzed in this study. This data can be found here: **The six discovery datasets are available through the NCBI Gene Expression Omnibus (GEO): https://www.ncbi.nlm.nih.gov/geo/query/acc.cgi?acc=GSE163151; https://www.ncbi.nlm.nih.gov/geo/query/acc.cgi?acc=GSE217948; https://www.ncbi.nlm.nih.gov/geo/query/acc.cgi?acc=GSE161731; https://www.ncbi.nlm.nih.gov/geo/query/acc.cgi?acc=GSE107991; https://www.ncbi.nlm.nih.gov/geo/query/acc.cgi?acc=GSE107992; and https://www.ncbi.nlm.nih.gov/geo/query/acc.cgi?acc=GSE101705. The COVID-19 validation dataset is available through GEO at https://www.ncbi.nlm.nih.gov/geo/query/acc.cgi?acc=GSE234207. Pulmonary-TB validation data are available through the Dryad Digital Repository associated with Hai et al. (2024):** https://doi.org/10.5061/dryad.s4mw6m9gf (28). No new primary datasets were generated in this study.

